# Extinction rate has a complex and non-linear relationship with area

**DOI:** 10.1101/081489

**Authors:** Petr Keil, Juliano S. Cabral, Jonathan Chase, Ines S. Martins, Felix May, Henrique M. Pereira, Marten Winter

**Affiliations:** German Centre for Integrative Biodiversity Research (iDiv), Halle-Jena-Leipzig, Deutscher Platz 5e, 04103 Leipzig, Germany; Institute of Biology, Leipzig University, Johannisallee 21, 04103 Leipzig, Germany; Ecosystem Modeling, Center for Computational and Theoretical Biology (CCTB), University of Würzburg, Germany; Institute of Biology, Martin Luther University Halle-Wittenberg, Am Kirchtor 1, 06108 Halle (Saale), Germany; Institute of Computer Science, Martin-Luther University Halle-Wittenberg, 06099 Halle (Saale), Germany; Cátedra Infraestruturas de Portugal-Biodiversidade, CIBIO/InBIO, Universidade do Porto, Campus Agrário de Vairão, 4485-661 Vairão, Portugal

**Keywords:** Anthropocene, continental, grain, habitat loss, island biogeography, local, mass extinction, MAUP, metapopulation, occupancy, patch, resolution

## Abstract

**Aim.:** Biodiversity loss, measured as count of extinction events, is a key component of biodiversity change, and can significantly impact ecosystem services. However, estimation of the loss has focused mostly on per-species extinction rates measured over limited numbers of spatial scales, with no theory linking small-scale extirpations with global extinctions. Here we provide such link by introducing the relationship between area and per-species probability of extinction (PxAR) and between area and count of realized extinction events in that area (NxAR). We show theoretical and empirical forms of these relationships, and we discuss their role in perception and estimation of the current extinction crisis.

**Location:** USA, Europe, Czech Republic, Barro Colorado Island

**Methods:** We derived the expected forms of PxAR and NxAR from a range of theoretical frameworks based on theory of island biogeography, neutral models, and species-area relationships. We constructed PxAR and NxAR in five empirical datasets on butterflies, plants, trees and birds, collected over range of spatial scales.

**Results:** Both the theoretical arguments and empirical data support monotonically decreasing PxAR, i.e. per-species extinction probability decreasing with increasing area; however, we also report a rare theoretical possibility of locally increasing PxAR. In contrast, both theory and data revealed complex NxAR, i.e. counts of extinction events follow variety of relationships with area, including nonlinear unimodal, multimodal and U-shaped relationships, depending on region and taxon.

**Main conclusions:** The uncovered wealth of forms of NxAR can explain why biodiversity change (the net outcome of losses and gains) also appears scale-dependent. Furthermore, the complex scale dependence of PxAR and NxAR means that global extinctions indicate little about local extirpations, and vice versa. Hence, effort should be made to understand and report extinction crisis as a scale-dependent problem. In this effort, estimation of scaling relationships such as PxAR and NxAR should be central.

## INTRODUCTION

Biodiversity loss as a result of species extinctions is potentially one of the most serious environmental problems we face, and estimates of its rate and magnitude are needed for informed decisions and conservation policy. The Aichi target 12 under the Strategic Plan for Biodiversity 2011-2020 (www.cbd.int/sp/targets/) aims at preventing extinctions of known threatened species, and all regional and the upcoming global IPBES assessments (http://www.ipbes.net/) are committed to report past, present, and future trends of biodiversity. Given the importance of assessing extinction rates, it is striking that there are major unresolved issues related to its spatial scaling and metrics.

So far, large-scale extinction science has focused on estimation of per-species extinction rates, measured as number of extinctions per million species-years (E/MSY) (Barnosky *et al.*, 2011; Proenca & Pereira, 2013; Pimm *et al.*, 2014). Similar metrics have been used in island biogeography (Wu & Vankat, 1995) and metapopulation biology (Hanski, 1991), where they are termed per-species extinction rate or per-species extinction probability. All these metrics are independent on absolute number of species (*S*), which makes them comparable across epochs, regions, and taxa.

However, apart from the per-species rates, absolute counts of extinction events per unit of time (hereafter *N*_*X*_) should also be of major interest, since they affect species richness *S* – a quantity that is at the core of basic biodiversity science (Gaston, 2000) and which has been associated with ecosystem services (Hooper *et al.*, 2005; Cardinale *et al.*, 2012). Recently, a number of authors have been particularly interested in understanding how *S* changes through time (hereafter *ΔS*), with debate as to whether it is declining at all scales, or instead has more variable outcomes (Vellend *et al.*, 2013; Dornelas *et al.*, 2014; McGill *et al.*, 2015). However, compared to the per-species extinction rates, *N*_*X*_ has rarely been calculated [but see Tedesco et al. (2013)].

The role of spatial scale is a fundamental, but often overlooked, aspect of understanding patterns of extinct rates. Current extinction rates have been estimated at global and continental extents (Barnosky *et al.*, 2011; Alroy, 2015), and global extinctions are indeed crucial since they are irreversible. Yet, any global extinction is preceded by a series of local and regional extinctions (a.k.a. *extirpations*), which often have ecological or iconic significance, or both.

For example, although Danube Clouded Yellow (*Colias myrmidone*, Esper 1780) still survives in some parts of Europe, the butterfly was extirpated from the Czech Republic in 2006, and this local extinction event triggered considerable attention for its implications for landscape management across the whole continent (Konvicka *et al.*, 2007). More prominent examples include extirpations of the bison (Isenberg, 2001) or lion (*Panthera leo*, Linnaeus 1758) (Riggio *et al.*, 2012) – local loss of such keystone species has direct impact on local ecosystem services, while country-wide extirpations matter because countries are obliged to protect their species either through international treaties (e.g. Aichi targets) or through national legislations. Clearly, focusing solely at the global extinctions can underestimate extirpations at smaller scales, and vice versa, and hence there is a need to assess the current extinction crisis at multiple spatial scales simultaneously.

In this Concept paper we propose a new way of looking at extinction rates to address the issues of metric and scale of extinction. Specifically, we propose to jointly consider how both the per-species extinction rates *P*_*X*_ and the counts of extinctions *N*_*X*_ scale with area over which they are observed. We first provide several theoretical expectations for the scaling of *P*_*X*_ and *N*_*X*_ derived from the theory of island biogeography, from neutral theory of biodiversity, and from a simple model based on species-area relationship. We then demonstrate the scaling using five empirical datasets, covering local, regional and continental scales. We show that while *P*_*X*_ mostly decreases with area, *N*_*X*_ follows complex relationships with area; the key finding is that *N*_*X*_ in small areas can be lower, but also higher, than *N*_*X*_ in large areas, making it impossible to get the complete picture of the current extinction crisis from looking at the global extinctions only.

## TERMS AND DEFINITIONS

We classify an event as *extinction* when a species disappears in the focal area (present at time 1, but absent at time 2), regardless to whether it still survives outside of the area. When the focal area is the entire world, an extinction event becomes a *global extinction*. Thus, a local or regional extinction event may also be called local or regional *extirpation*, if the species still prevails outside of that locality or region.

We use *P*_*X*_ for *mean per-species probability of extinction* in a given area *A*, and *N*_*X*_ for *mean number of extinctions* in *A* during a specified time period. We use *mean* because we consider *P*_*X*_ and *N*_*X*_ as means of probability distributions over multiple species that occur in a given area. Thus, our *N*_*X*_ is similar to *E*/*t* metric described by Proenca & Pereira (2013), and our *P*_*X*_ is similar to their (*E*/*S*)/*t* metric. We use PxAR to describe the relationship between *P*_*X*_ and *A*, and we use NxAR to describe the relationship between *N*_*X*_ and *A*. We shall note here that PxAR has been referred to as an *extinction-area relationship* in Hugueny et al. (2011). This term is also used by Kitzes & Harte (2014) for a different concept: the probability of extinction conditional on the area of lost habitat (instead of the size of the sampled area). To prevent confusion, we avoid the term extinction-area relationship altogether.

We use *spatial scale* as a synonym for *sampling area, resolution, and/or grain*; that is, the area (*A*) of an observation window (grid cell, country) in which extinction events are counted. We focus exclusively on spatial scaling, disregarding the role of *temporal scale* (i.e., the length of temporal window over which extinctions are counted (Barnosky *et al.*, 2011). We circumvent the problem by assuming the temporal window approximately or exactly constant in each individual dataset or in a particular theoretical argument. However, the temporal scaling of extinction rates is a critical issue that should be addressed in future research.

Some existing work on spatial scaling of extinction rates is rooted in island biogeography and metapopulation theory, where extinction rates are calculated over non-nested islands or habitat patches of varying size (Scheiner, 2003). We call these *island systems*. In contrast, almost no attention has been focused on spatial scaling of extinction rates in continuous regions, in which a region is overlaid by a system of nested grids of increasing resolution (Storch *et al.*, 2012; Keil *et al.*, 2015), and *P*_*X*_ and *N*_*X*_ are calculated at each resolution. Grid cells in the grid can be either regular rectangles, or irregular administrative divisions, but are always aggregated (up-scaled) in a strictly nested way, and hence we call them *nested systems* – as such they are similar to the Type I species-area curves of (Scheiner, 2003).

To avoid confusion, we note that our treatment of spatial scaling of extinction rates is fundamentally different from approaches using the *Endemics-Area Relationship* (EAR) and backward *Species-Area Relationship* (SAR) to predict extinction rates due to habitable area loss (He & Hubbell, 2011; Pimm *et al.*, 2014; Keil *et al.*, 2015) (Fig. 1). The EAR-describes a hypothetical impact one specific driver of extinction; loss of habitable area. In contrast, PxAR and NxAR provide spatial scaling of a *dynamic* process of the actual realized extinctions in a given area of observational window *A* and over given temporal window, without regard to the causes of extinction (habitat loss, global warming, pathogens, hunting, or invasions; see Fig. 1 for details).

**Figure 1.**
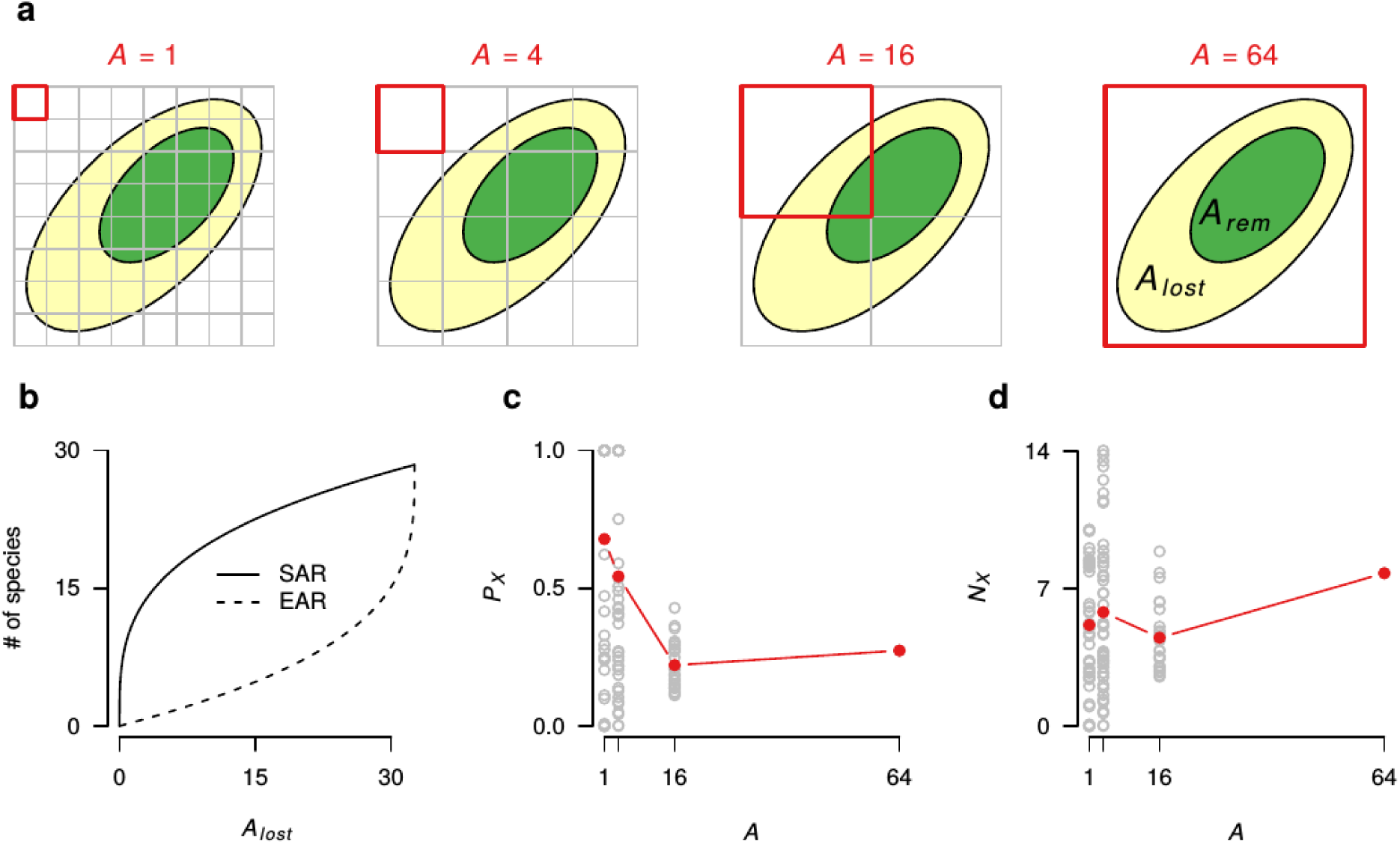
Difference between spatial scaling of realized extinction rates (PxAR and NxAR) in nested systems and static scaling of species richness (SAR, EAR) commonly used to estimate extinction rates due to habitat loss (Pimm & Raven, 2000; He & Hubbell, 2011; Keil *et al.*, 2015). (a) Hypothetical landscape where part of the habitat area is lost (*A*_*lost*_; pale yellow), and part remains (*A*_*rem*_; green). (b) Number of species extinct in *A*_*lost*_ is described by the endemics-area relationship (EAR). Since we assume that individuals in the habitat are distributed randomly, EAR follows a point reflection symmetry with species-area relationship (SAR) (Keil *et al.*, 2015); here SAR takes the form of *S* = 10*A*^0.3^. In contrast to EAR, PxAR (c) and NxAR (d) are calculated over the observation window (red) and over a given temporal period. Dots in panels c and d are values calculated directly using EAR described in panel b. Red dots correspond to the mean PxAR and mean NxAR observed across all windows of a given temporal period.

## THEORETICAL EXPECTATIONS

We offer several complementary ways of reasoning, showing that *N*_*X*_ and *P*_*X*_ should be expected to follow a wealth of functional responses to *A*. These are not meant as theories to be tested per se; rather, they are different ways to think about spatial scaling of extinctions in different systems and over different spatial grains and extents.

### Scaling of per-species extinction probability *P*_*X*_

Here we show that, theoretically, the relationship between *P*_*X*_ and *A* should mostly be negative, with some exceptions.

#### Island systems

There is a general expectation that per-species probability of extinction (*P*_*X*_) decreases with area of an island. This is supported both empirically (Diamond, 1984; Quinn & Hastings, 1987; Hugueny *et al.*, 2011) and theoretically (Hanski & Ovaskainen, 2000). The paradigm is that larger areas can host larger populations, which are less likely to perish due to environmental and demographic stochasticity (Lande, 1993). The latter mechanism is implicitly assumed in the equilibrium theory of island biogeography (ETIB) (MacArthur & Wilson, 1967) and in spatially realistic models in metapopulation ecology (Hanski & Ovaskainen, 2000; Hanski, 2001). An example of a specific scaling function assumed in metapopulation ecology is a hyperbola *P*_*X*_ = *e*/*A*, where *A* is a patch area and *e* is a constant reflecting both population density in a unit of area and rate of reproduction per unit of time (Hanski & Ovaskainen, 2000), with the assumption that extinction risk scales as an inverse of population size in taxa affected by moderate environmental stochasticity (Lande, 1993), and population size is proportional to area of habitat patch or island.

#### Neutral models

The mentioned hyperbolically decreasing PxAR has also been described in neutral models (Ricklefs, 2006; Halley & Iwasa, 2011). However, Ricklefs (2006) only examined extinction rates at the scale of the entire metacommunity, while Halley & Iwasa (2011) disregarded the potentially important role of immigration, and they generated their abundance distributions by ad-hoc mechanisms, not by the neutral model itself. To get a hint of possible forms of PxAR in neutrally modeled local communities with immigration, we simulated from a neutral model presented by Hubbell (2001), which we parametrized using the classical combination of parameters used in Hubbell (2001) (page 135): *Θ*; = 50, *J* = 5 × 10^5^, local mortality rate *D* = 0.1 and immigration rate *m* = 0.1. We simulated 50 local communities, each with 10, 100, 1,000 and 10,000 individuals, which we converted to area [ha] assuming the density of 24,341 individuals per 50 ha (Hubbell, 2001). We initialized the local community with a single species, discarded the first 1,000 simulation steps as burn-in, and looked at *P*_*X*_ after 5 generations and also after 50 generations (Fig. 2).

**Figure 2.**
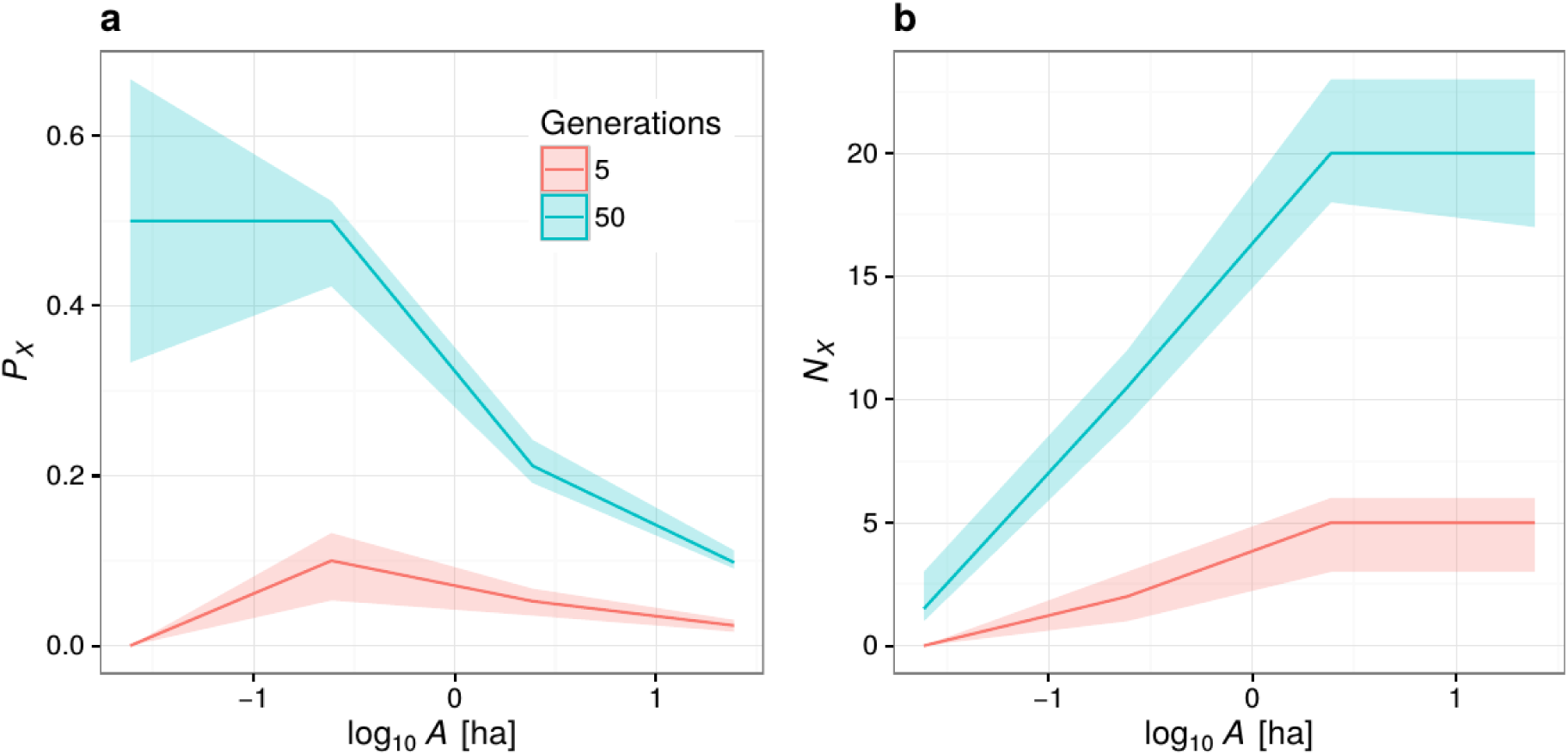
PxAR and NxAR from simulated realizations of the classic version of the spatially implicit neutral model (Hubbell, 2001). The solid lines and ribbons show medians and quartiles of the simulations.

We found that *P*_*X*_ decreased with area of the local community after 50 generations, but the decrease was initially shallower than expected in a hyperbolic decay proposed by Ricklefs (2006) and Halley & Iwasa (2011) (Fig. 2a). This suggest the role of immigration in reducing local *P*_*X*_, since immigration is what is missing in the neutral drift models of Ricklefs (2006) and Halley & Iwasa (2011). Surprisingly, the PxAR was hump-shaped when *P*_*X*_ was measured over only 5 generations (Fig. 2a), highlighting the crucial role of temporal scale, which we otherwise ignore in this paper.

#### Heterogeneous nested non-equilibrium systems

The mentioned assumption of proportional relationship between population size and area is reasonable when focusing solely on area of homogeneous patches of habitats with randomly or regularly distributed populations within habitats. However, the assumption falls short in nested systems, where observation windows contain multiple habitats, where populations are spatially aggregated, and where the magnitude of the aggregation varies with scale. In such situation we can express *P*_*X*_ at a given scale as the proportional loss of occupied grid cells of area *A* (Fig. 3a) – an idea that builds on the framework outlined by Kunin (1998) and later applied to temporal dynamics of species distributions by Wilson et al. (2004). Our novel point here is that *P*_*X*_ calculated in nested systems can decrease, but also *increase* with *A*, depending on the range of *A* values that is examined, and depending on where exactly are populations lost (see Fig. 3a for full exposition).

**Figure 3.**
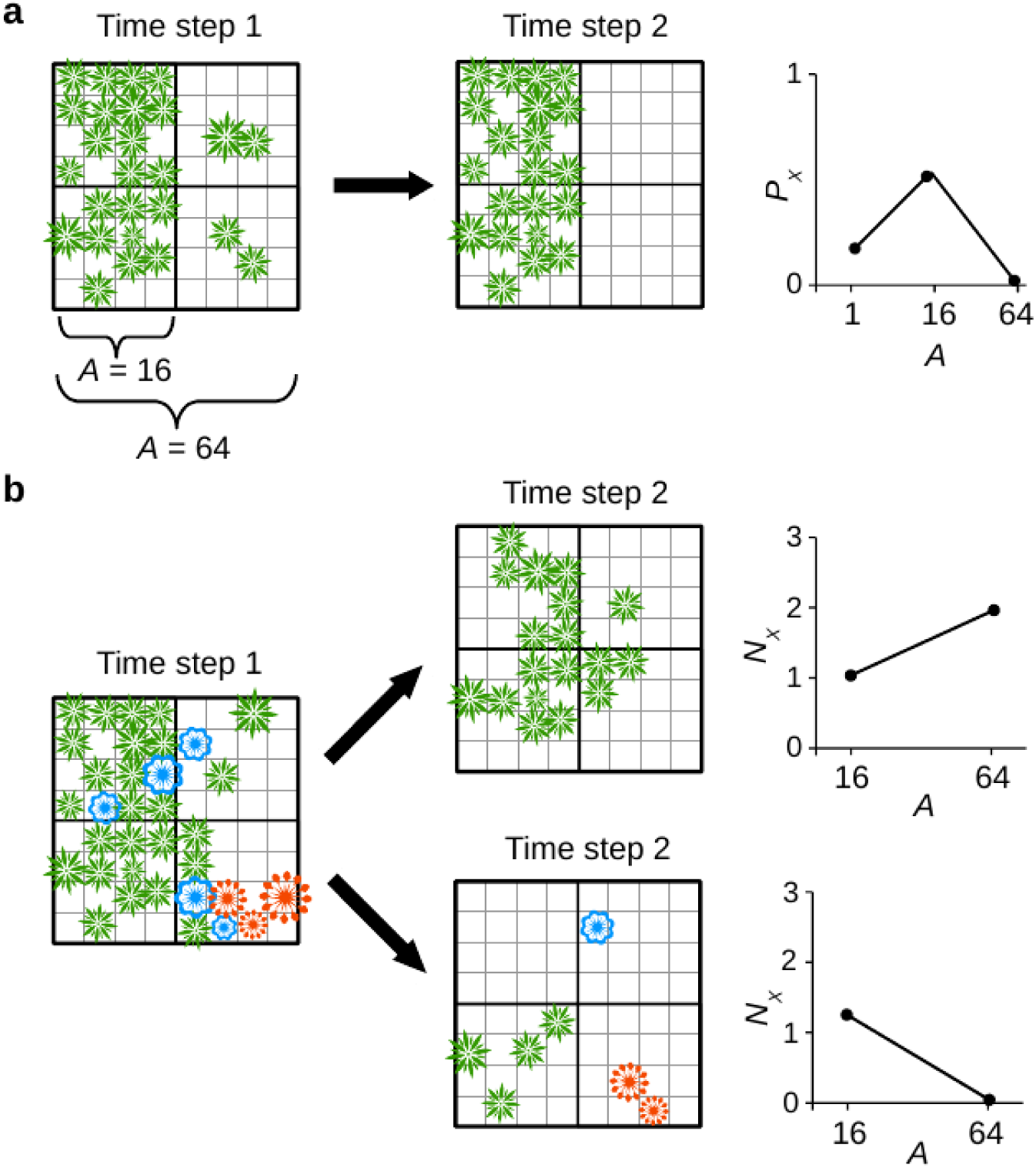
Simple scenarios producing both increasing and decreasing shapes of PxAR (a) and NxAR (b) in nested systems. In panel (a) we demonstrate the theoretical possibility of *P*_*X*_ increasing with area *A*. In panel (b) we show two kinds of community dynamics producing opposite directions of NxAR in a plant community that started with 3 species (blue, green, red) at time step 1. In the first scenario, two rare species go extinct, leading to increasing NxAR. In the second scenario, none of the species goes completely extinct, but there are severe population declines in all species, leading to decreasing NxAR. Values of *P*_*X*_ and *N*_*X*_ in the graphs are averages over all four areas with *A* = 1 and *A* = 16.

### Scaling of counts of extinctions *N*_*X*_

Here we show that the relationship between *N*_*X*_ and *A* can follow wealth of functional forms, including negative, positive, or non-linear.

#### Island systems

The equilibrium theory of island biogeography (MacArthur & Wilson, 1967) can be used to derive the NxAR. The ETIB-based model considers equilibrium extinction rate, i.e. extinctions are in balance with species gains. According to ETIB, the equilibrium rate *N*_*X*_ is:

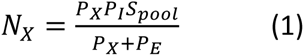

Where *S*_*pool*_ is number of species in species pool, *P*_*I*_ is per-species probability of immigration, and *P*_*E*_ is per-species probability of extinction (Wu & Vankat, 1995). We assume that *P*_*X*_ = 0.01^/^*A* (the hyperbola mentioned above). Critical is the relationship between *P*_*I*_ and *A*, which we present in two forms: (1) *P*_*I*_ *is constant* (Fig. 4a) and independent on *A*. In the simplest formulation of ETIB, there is no relationship between *P*_*I*_ and *A*. The resulting NxAR in this case is always monotonically decreasing (Fig. 4a). (2) *P*_*I*_ *increases with area.* In this scenario, increasing area means lower *P*_*I*_ as before, but also higher immigration rates because larger areas are larger targets (Lomolino *et al.*, 2010); hence, we introduce a positive linear proportional relationship between *P*_*I*_ and *A*. This generates a hump-shaped NxAR (Fig. 4b).

**Figure 4.**
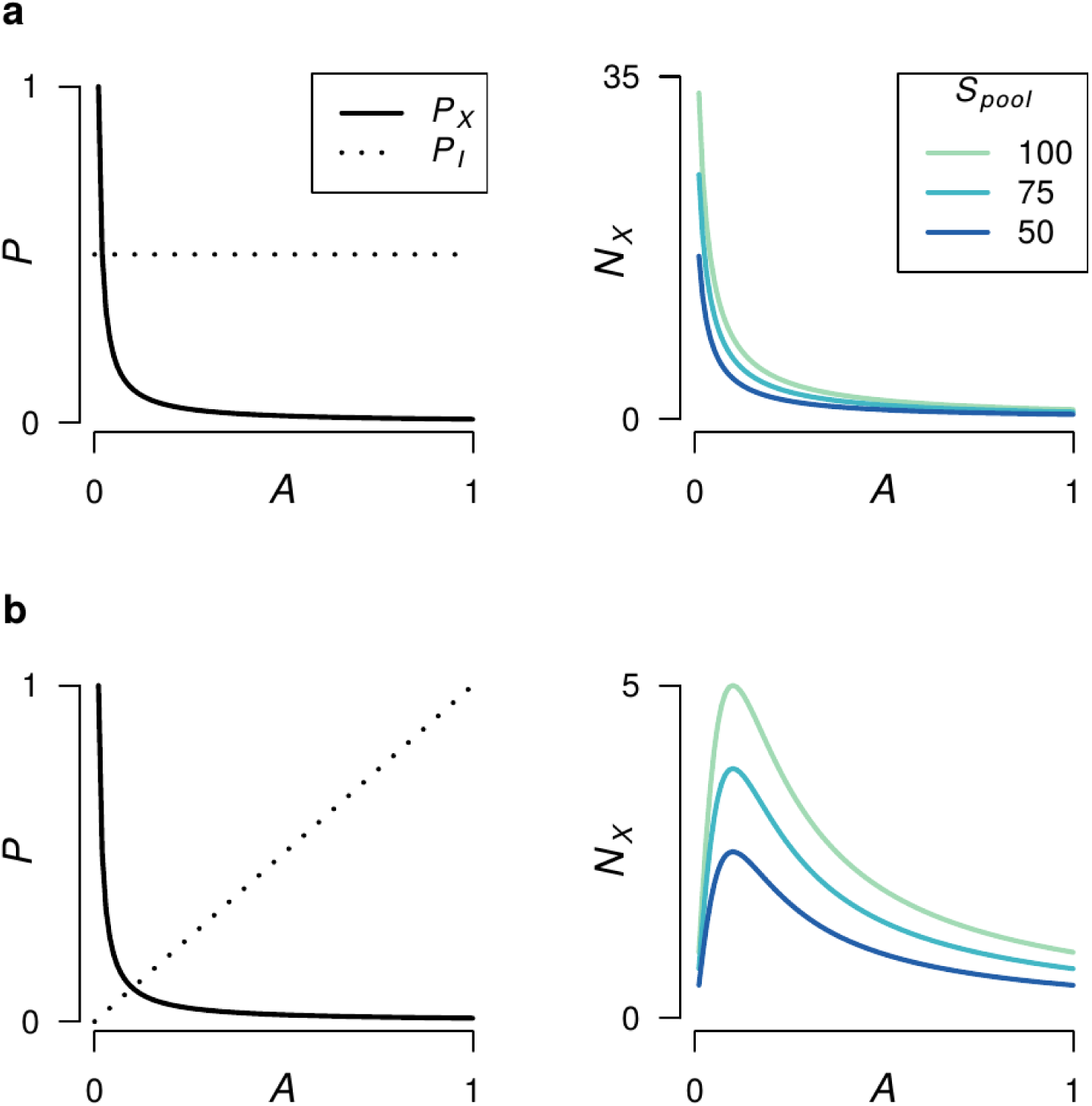
Two scenarios describing spatial scaling of per-species extinction probability *P*_*X*_, per-species immigration probability *P*_*I*_, and the resulting relationship (NxAR) between area *A* and number of extinctions *N*_*X*_ in island systems. (a) *P*_*X*_ decreases proportionally with area, *P*_*I*_ is constant. (b) *P*_*X*_ decreases with area, *P*_*I*_ increases with area. The NxAR curves were calculated using eq. 1.

Other NxAR shapes are likely to emerge from different relationships between *A* and *P*_*X*_, *P*_*I*_ and *S*_*pool*_. Additionally, limiting the range of areas that are being observed (x-axis of the NxAR plot) can also lead to a monotonically increasing or decreasing NxAR. Therefore, in the ETIB model, various shapes of NxAR are produced by the interplay between per-species extinction probability with per-species immigration and the size of the species pool.

#### Neutral models

We are not aware of any literature examining *NxAR* in neutral models. Our parametrization of the spatially implicit neutral model [see Hubbell (2001) and the previous section] predicts positive relationship between *N*_*X*_ and area of local community, measured over the limited range of spatial and temporal scales that we examined (Fig. 2b).

#### Heterogeneous nested non-equilibrium systems

Here we show two lines of reasoning, both leading to a wealth of NxAR shapes.

First, let us consider a nested system and a simple verbal qualitative model. Imagine two scenarios that lead to different NxARs (Fig. 3b): (1) A continent loses several geographically extremely restricted species, and nothing happens to the widespread species. The localized extinctions will have almost no effect on *N*_*X*_ at small scale, but will increase *N*_*X*_ at the continental scale. In this scenario, *N*_*X*_ *increases* with *A*. (2) There are range contractions in several widespread species, but no species goes completely lost from the continent. This results in high *N*_*X*_ at small scale, but no extinctions at the continental scale. Here, *N*_*X*_ *decreases* with *A* (Fig. 3b).

In the second line of reasoning, we can assume that mean species richness *S* at each grain follows the *species-area relationship* (SAR), for example a power law (Fig. 5a). We then invoke a one-time ad-hoc stress event during which each species at each location either goes extinct or survives. We assume that, on average, the per-species extinction probability (*P*_*E*_) monotonically decreases with *A*, but that monotonic decreases can have various forms (Fig. 5b). In such system, NxAR is obtained by multiplication of SAR with PxAR, or formally: If *S* = *f*(*A*) and *P*_*X*_ = *g*(*A*), then number of extinctions (*N*_*X*_) per area of observation window (*A*) is *N*_*x*_ = *f*(*A*)*g*(*A*). As a result, the emerging shapes of NxAR can vary from increasing to hump-shaped (Fig. 5c), with the possibility of observing monotonically decreasing NxAR over limited extents of *A* (Fig. 5c).

**Figure 5.**
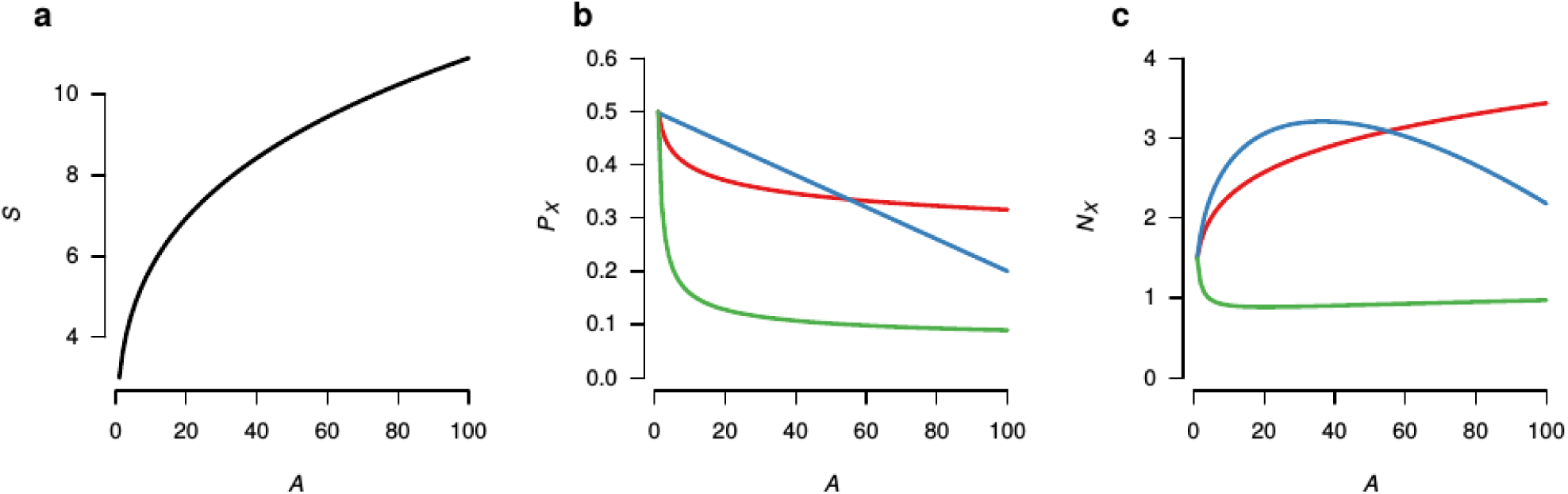
Hypothetical examples of spatial scaling of number of extinction events (*N*_*X*_) derived from a simple SAR-based model. (a) Species richness *S* follows a power-law with area of observation window *A*. (b) Per-species extinction probability *P*_*X*_ follows various monotonically decreasing scaling relationships with *A*, which results in a rich variety of NxARs (c). Colors indicate corresponding PxARs and NxARs.

## EMPIRICAL EVALUATION

We calculated empirical NxAR and PxAR using five datasets, with the aim to cover wide range of geographic extents and types of temporal dynamics. We focus on terrestrial mainland nested systems, mostly due to our own research background, and because there already is some literature on empirical extinction rates in island-like systems (Diamond, 1984; Quinn & Hastings, 1987; Hugueny *et al.*, 2011; Tedesco *et al.*, 2013), although the literature does not cover NxAR. Hence, an assessment of empirical PxAR and NxAR on islands is an important future task.

### Data

#### European butterflies

We extracted data on extinction events and on the extant species of *butterflies* (Lepidoptera: Rhopalocera) in European administrative areas during the last ~ 100 years from Red data book of European butterflies (Van Swaay & Warren, 1999); specifically, we considered species extinct in a given country if its status in Van Swaay & Warren (1999) Appendix 6 table was “Ex”.

#### European plants

We extracted the same kind of data on extinctions (between AD 1500 and 2009) and extant species of European *vascular plants* from Winter et al. (2009), available upon request from MW. Note that the plant data cover different extent and administrative units than the butterfly data (Fig. 6).

**Figure 6.**
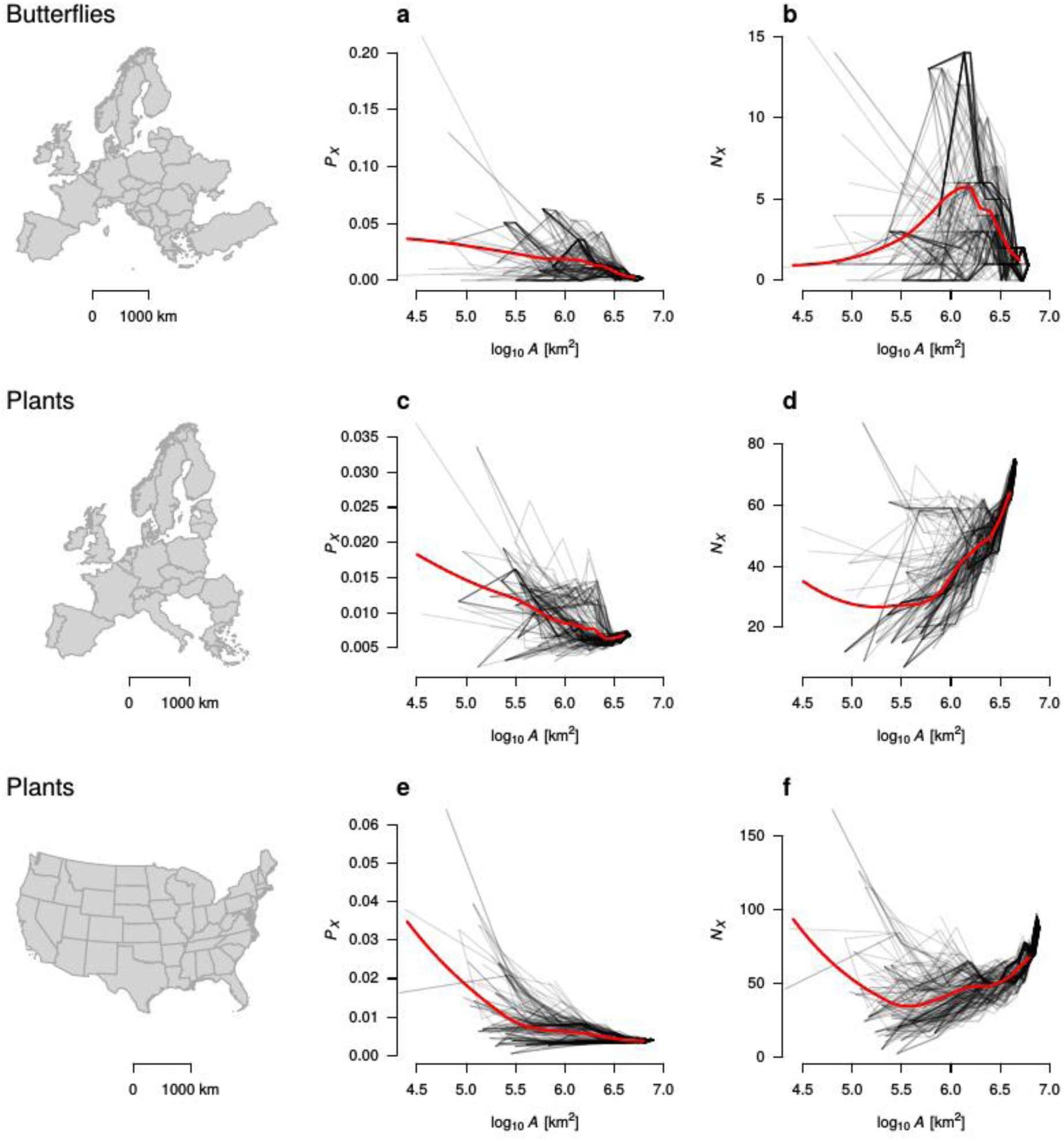
Empirical spatial scaling of per-species probability of extinction *P*_*X*_ (a, c) and number of extinctions *N*_*X*_ (b, d) of European butterflies (a, b) and European vascular plants (c, d) and US vascular plants (e, f). Each grey line was obtained by placing (200 times) a small circle at a random location within Europe, then gradually increasing the circle size up to the size of the whole continent, counting number of extinction events within the countries falling in the circle. Red lines are LOESS regressions with smoothing span of 0.3.

#### United States plants

We used data on the extant native species of *Plantae* (ferns, conifers and flowering plants) in 47 states of the USA (i.e. excluding Hawaii, Alaska, and Washington DC) provided in the BONAP’s North American Plant Atlas (Kartesz, 1999); for an example use of these data in a quantitative analysis see Winter et al. (2010). We used data on contemporary (AD 1500 - 2009) extinction events also at the level of individual states. This information is provided by NatureServe (www.natureserve.org) and its network of Natural Heritage member programs, a leading source of information about rare and endangered species, and threatened ecosystems (NatureServe, 2016). The data provided by NatureServe are for informational purposes, and should not be considered a definitive statement on the presence or absence of biological elements at any given location. Site-specific projects or activities should be reviewed for potential environmental impacts with appropriate regulatory agencies.

#### Czech birds

We used digitized presences and absences of breeding birds recorded in a regular grid of 11.7 x 11.7 km grid cells in the Czech Republic, and covering two temporal periods: 1985-1989 (Šťastný *et al.*, 1997) with 56,780 species-per-cell incidences, and 2001-2003 (Šťastný *et al.*, 2006) with 59,354 species-per-cell incidences.

#### Barro Colorado 50ha plot

We used tree data from 50-ha forest plot from Barro Colorado Island (BCI), Panama (Condit, 1998; Hubbell *et al.*, 1999, 2005). We only included trees with ≥ 10 cm DBH, and we compared two temporal snapshots that are 25 years apart. The results were qualitatively similar when the temporal lag between the two snapshots was 5, 10, 15 and 20 years, and also when we considered all trees with ≥ 1 cm DBH.

### PxAR and NxAR calculation

In the European, US and Czech datasets, we constructed PxAR and NxAR curves by placing a small circle at a random location within the geographic extent of the data, then gradually increasing the circle size. For each circle size, we aggregated spatial units (EU countries, US states, CZ grid cells) overlapped by the circle. In the resulting aggregated units we noted *A*, *P*_*X*_, *N*_*X*_ and *S*, where *A* is the area of the aggregated unit, *N*_*X*_ the number of extinction events, *S* the number of all species found in the aggregated unit in the first temporal window, and 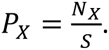. This whole procedure was repeated 200 times, each time starting at a different random location of the smallest circle. We obtained a set of curves for each starting position, which we summarized using LOESS regression with smoothing span of 0.3.

In the BCI forest plot, we overlaid the entire plot area with rectangular grids of increasing resolution, from 20 m × 20 m up the the entire 50 ha plot. In each grid cell we counted the number of species (*S*_0_) in the 1985 census and checked which of these species were present in the 2010 census (*S*_1_). We then calculated *P*_*X*_ = ^(^*S*_0_ − *S*_1_^)⁄^*S*_0_ and *N*_*X*_ = *S*_0_ − *S*_1_. We calculated average (± standard deviation) *P*_*X*_ and *N*_*X*_ over all grid cells at a particular grid resolution.

### Results

In the five empirical datasets, *P*_*X*_ decreases with area *A* of observation window (Figs. 6a, c, e, 7a, c). This contrasts with *N*_*X*_, which scales with *A* in a variety of ways. The two datasets from Europe, plants and butterflies, produced highly divergent NxARs (Fig. 6): European butterflies (Fig. 6b) had a hump-shaped NxAR, with only a single pan-continental extinction, but with pronounced losses at intermediate scales. In contrast, European vascular plants (Fig. 6d) have an upward-accelerating NxAR, with highest rates of extinction at the largest scale. NxAR of the US plants is similar to the NxAR of the European plants (Fig. 6f), with clear upward-acceleration at large areas.

A decreasing NxAR is found in Czech birds (Fig. 7b). Here we note that the high values of *N*_*X*_ at small scales can be a result of under-sampled grid cells. On the even smaller scale of the 50 haBCI forest plot (Fig. 7d), the NxAR first sharply increases at the finest scale, with humps at larger scales, unveiling the possibility of multi-modal empirical ExARs.

**Figure 7.**
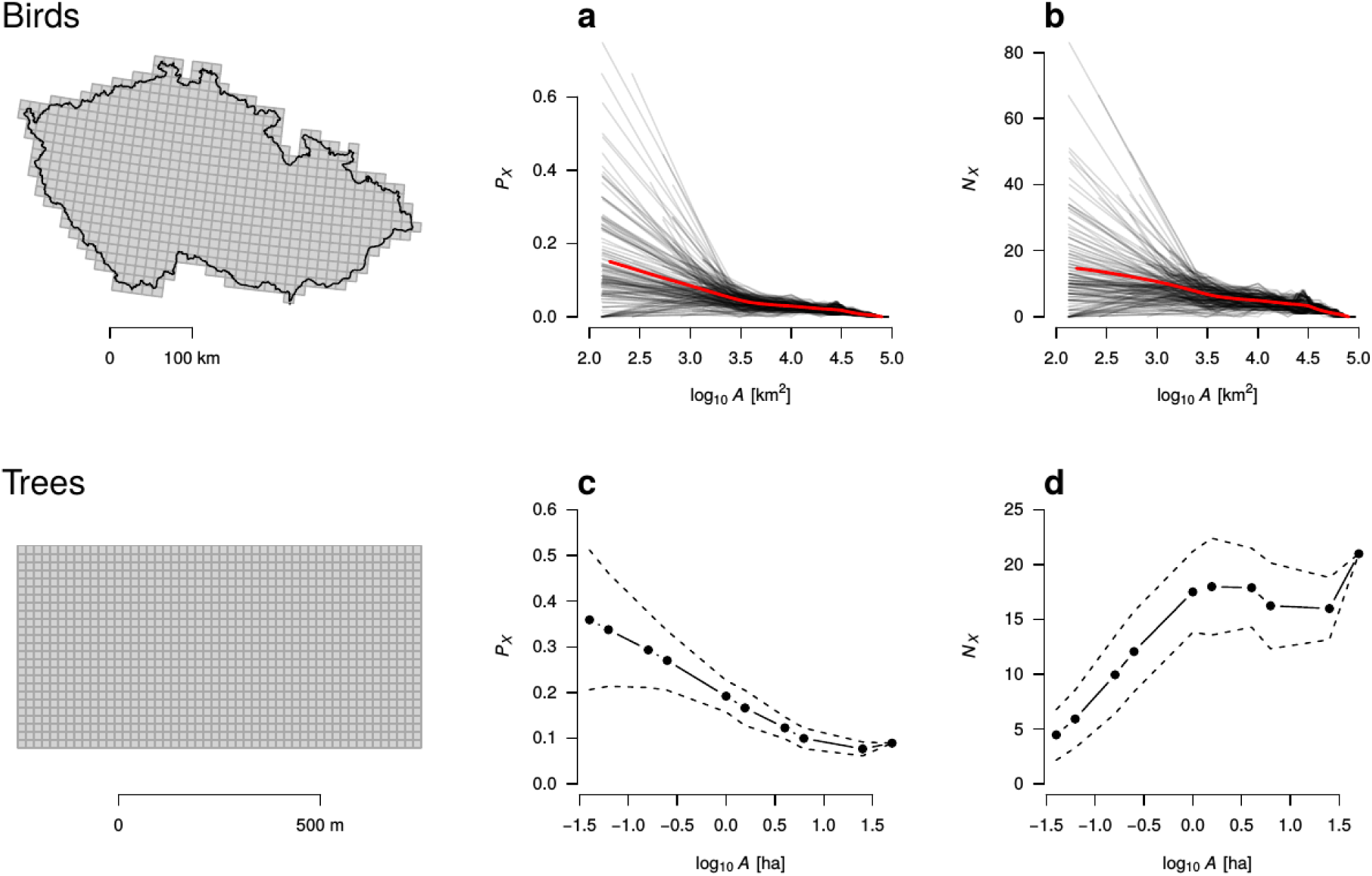
Empirical spatial scaling of per-species probability of extinction *P*_*X*_ (a, c) and number of extinctions *N*_*X*_ (b, d) in a country-wide and a local dataset. Panels a and b use atlas data on birds of Czech Republic (a, b). Each grey line was obtained by placing (200 times) a small circle at a random location within the Czech Republic, then gradually increasing the circle size, counting numbers of extinctions in atlas cells overlapping the circle. Red lines are LOESS regressions (span = 0.3). High values of *P*_*X*_ and *N*_*X*_ in some small Areas are likely caused by undersampling. Panels c and d use data on trees with DBH ≥ 10 cm in the 50 ha BCI forest plot (c, d) calculated over a 25-year lag. Solid circles and lines are means, dashed lines are standard deviations calculated over different spatial locations of the observation window.

## DISCUSSION

We have demonstrated both theoretically and empirically that extinction rates do not follow a simple monotonic relationship with area. Instead, both local and regional extinction rates can be low, with high rates at intermediate regional scales – or the situation can be completely the opposite. Moreover, even curved relationships can be expected.

The observations of negative PxAR and positive NxAR at the local scale of BCI plot are in line with predictions of the neutral model. In both cases *N*_*X*_ approaches 0 as *A* approaches 0, which stems from the trivial fact that limited number of individuals at small areas can only be divided in a limited number of species. Interestingly, as *A* increases and approaches the size of the BCI plot, both the neutral model and the empirical BCI data reveal a flat phase of the NxAR; we suggest that this is because the strong limiting effect of small number of individuals no longer applies above a certain *A*. Future research should focus on more realistic versions of the neutral model, perhaps in a spatially explicit setting (Rosindell & Cornell, 2007), and over larger spatial extents.

In contrast with the BCI plot, it is unreasonable to assume equilibrium or neutral dynamics at the extent of the entire countries and continents, especially when considering contemporary human-induced extinctions and extirpations. In such situations it is more instrumental to explain the observed extinction scaling with range contractions of rare and widespread species (see the Theoretical expectations section): The observed *decrease* of *N*_*X*_ with *A* at large scales in European butterflies likely reflects range contractions, but not complete extinctions, of widespread species. This is the opposite of the observed *increase* of *N*_*X*_ with *A* at large scales in European and US plants, which reflects complete losses of small-ranged species. Importantly, these disparate directions of NxAR are perfectly reconcilable with the decreasing PxAR in all of these datasets, as we have described in the SAR-based model (see the Theoretical expectations section and Figure 5).

For the first time, we have shown that in theory, *per-species probability of extinction* can increase with area. Seemingly, this goes against the classical views (MacArthur & Wilson, 1967; Diamond, 1984; Quinn & Hastings, 1987; Hanski, 2001) – how can a species be more likely to go extinct in a large area than in small one? One key is a potential existence of remote populations occupying a small fraction of an entire region (i.e., a country). Loss of such small population can mean a complete extinction in the entire country, but may not contribute to the small-scale *P*_*X*_, since the species still retains numerous small-scale populations outside of the country. Is this an artifact of the specific position of the observation windows? Probably not, since some parametrizations of the neutral model can also give a higher *P*_*X*_, and since the model is spatially implicit (the observation window has size, but no position). Note: In our empirical data, we detected only negative PxAR, suggesting that it is still the prevalent form in real-world nested systems.

The uncovered wealth of possible relationships of *counts of extinctions* with area has several crucial implications. First, it can reconcile the reported lack of biodiversity change, on average, at local scales (Vellend *et al.*, 2013; Dornelas *et al.*, 2014) with reported losses at global scales (Barnosky *et al.*, 2011; Alroy, 2015). We are aware that, for a complete reconciliation, one may also need to know the spatial scaling of species gains (Jackson & Sax, 2010), which we see as an important future research avenue. However, the important lesson here is that even when species gains are ignored, or when they are scale-invariant, NxAR alone is sufficient to cause scale-dependent biodiversity dynamics. Furthermore, our results on extinctions indicate that biodiversity change at intermediate regional scales can be completely different from both local and global patterns. Therefore, simple *interpolation* between scales should be done with caution, if at all.

This leads to implications of our results for *extrapolation* of extinction rates at grains for which we have limited data (He & Hubbell, 2011; Pimm *et al.*, 2014). Given the richness of possible NxAR shapes, we warn that estimating magnitude of global extinction crisis from local species loss can be challenging. Similarly, numbers of local extinctions will be tricky to predict from global or continental extinction numbers. We suggest that, in both cases, extra information is necessary on spatial scaling of the drivers of extinction (e.g. on spatial scaling of habitat dynamics or spatial scaling of intensity of extinction debts), on the specific distributions of species within the studied area, and on their theoretical links to NxAR.

An alternative, and perhaps more promising approach could be to extrapolate PxAR, since in our five datasets it was always decreasing and hence more predictable than NxAR. If in the future there is a broader empirical support for an exact form of PxAR, then it should be possible to combine it with SAR (Fig. 4) derived from the current global distributional information on some taxa (i.e. the IUCN range maps) to reliably extrapolate extinction counts to grains that are coarser or finer than the grain of the extinction data. This can be a viable alternative to the recent approaches that predict extinction numbers solely from habitat loss (Pimm & Raven, 2000; Pereira & Daily, 2006; He & Hubbell, 2011; Keil *et al.*, 2015). Further, a predictable PxAR can be instrumental in statistical models that link patterns of biodiversity with environmental drivers at multiple spatial scales (McInerny & Purves, 2011; Keil *et al.*, 2013; Keil & Jetz, 2013).

Although we have focused exclusively on extinction, we suggest that a complementary approach can be used to study species gains, which includes immigration and speciation. This could be an exciting new opportunity especially for invasion biology. Another fruitful research can involve a more detailed comparison of the nested and island systems. It is well known that island and nested (mainland) systems differ in their static scaling patterns of biodiversity, but the causes of such discrepancy remain controversial (Whittaker *et al.*, 2001; Triantis *et al.*, 2012). In this paper, we suggested that nested observation windows that contain multiple habitat types can potentially differ in their PxAR from island systems of homogeneous habitat patches. However, the differences may potentially be elsewhere, and may be related to the same mechanisms that cause the differences, e.g. between the island and mainland SARs (Pereira & Borda-de-Água, 2013).

To conclude, our results underscore the need to treat species loss as a scale-dependent problem, or more precisely, a grain-dependent problem. By focusing at global extinctions only, we can miss that either not much is happening at small grains, or that there are drastic losses at the small grains. To provide a complete picture, local, regional and global losses should be reported at the same time, simultaneously. Here the PxAR and NxAR curves offer a straightforward way to summarize and report such multi-scale species loss, and we argue that estimation of these relationships can benefit assessments of the past and current state of biodiversity, as well as in forecasts of biodiversity’s future.

## ACKNOWLEDGEMENTS

We are grateful to A. Berger and C. Meyer for helpful comments. The BCI forest dynamics research project was founded by S.P. Hubbell and R.B. Foster and is now managed by R. Condit, S. Lao, and R. Perez under the Center for Tropical Forest Science and the Smithsonian tropical Research in Panama. Numerous organizations have provided funding, principally the U.S. National Science Foundation, and hundreds of field workers have contributed.

## DATA ACCESSIBILITY

Digitized form of the European butterfly data is available on Figshare at https://dx.doi.org/10.6084/m9.figshare.4036236.v1. An electronic form of the Czech bird dataset is not publicly available; contact PK for details. Data on extant species of plants in US states and European countries, and on plant extinctions of European countries are also not publicly available; contact MW for details. Data on US plant extinctions were purchased from NatureServe. The BCI data are available from CTFS through a request form (http://ctfs.si.edu/webatlas/datasets/bci/).

## BIOSKETCH

**Petr Keil** is a postdoc at iDiv. His main interest is spatial scaling of multiple facets of biodiversity and its change. His hobby is statistics.

**German Centre for Integrative Biodiversity Research (iDiv)** is a research institute run by Martin Luther University Halle-Wittenberg (MLU), Friedrich Schiller University Jena (FSU) and Leipzig University (UL) – and in cooperation with the Helmholtz Centre for Environmental Research (UFZ). The goals of the institute, which was set up back in 2012, are to record biological diversity in its complexity, to make available and use scientific data on a global level, and to develop strategies, solutions and utilization concepts for decision-makers in order to prevent any further loss of biodiversity. iDiv promotes theory-driven synthesis, experiments and data-driven theory in biodiversity sciences, and provides scientific foundation for sustainable management of Earth’s biodiversity.

